# Wounding induces preexisting multinucleated cells to survive and thrive as leader cells

**DOI:** 10.1101/2025.06.24.661427

**Authors:** Yuta Takahashi, Kenji Matsuzawa, Junichi Ikenouchi

## Abstract

Epithelial wounds are repaired through collective cell migration, a process orchestrated by a small subset of leader cells at the wound edge^1–3^. How these functionally distinct cells arise from an apparently homogeneous population of epithelial cells remains unclear. Here, we show that injury to cultured epithelial sheets allows the survival of multinucleated cells that are otherwise eliminated under normal conditions. We reveal that multinucleated cells preexist prior to injury and extend protrusions toward the wound, eventually adopting leader-like behaviors. These findings identify multinucleated cells as a latent reservoir for leader cell emergence. Our work highlights the inherent heterogeneity of epithelial sheets and uncovers a previously unrecognized function of multinucleated cells during wound healing.

## Introduction

Injury to epithelial sheets compromises their barrier function, allowing pathogens and foreign substances to enter the body from the external environment, while also causing the loss of fluids and nutrients from within, both of which pose a serious threat to the survival of the organism ^4–6^. Accordingly, large-scale wound to epithelial tissues must be repaired rapidly and efficiently through a process known as wound healing^7–9^.

During epithelial wound healing, cells coordinate their movement to quickly repopulate and cover the injured area. A key player in this collective migration is the “leader cell”, which enhances the motility of the epithelial sheet by exerting traction forces on surrounding follower cells^3,10^. A widely accepted model for the emergence of leader cells posits that cells located at the wound edge differentiate into leader cells in response to wound-derived signals in a process called wound-induced specification^11,12^. However, this model does not fully explain why only a small subset of wound-adjacent cells adopt leader cell fates, despite all these cells being similarly exposed to injury cues.

In this study, we focused on the observation that many leader cells are multinucleated and hypothesized that such cells, preexisting within the epithelial sheet, serve as a latent reservoir for leader cell differentiation upon tissue injury. Multinucleated cells are typically recognized as abnormal, pre-malignant cells and are rapidly eliminated from the epithelium under homeostatic conditions^13–16^. However, we found that epithelial injury suppresses this elimination in an Akt activation-dependent manner, thereby prolonging the survival of these cells. Our findings reveal a new model of leader cell emergence that expands the current understanding by incorporating the preexisting cellular heterogeneity and the dynamic balance between the elimination and persistence of multinucleated cells in epithelial tissues.

## Results

### Multinucleated cells are sporadically found within epithelial sheets and are often associated with elevated p53 activity

Previous studies have shown that defects in cytokinesis, frequently triggered by mechanical imbalances such as uneven tensile stress across the monolayer, can lead to accidental multinucleation ^17–19^. To determine the prevalence of multinucleated cells in epithelial monolayers, we used MDCKII cells, a canine kidney-derived epithelial cell line, and analyzed the cellular composition of confluent monolayers. Immunostaining of fixed cells with antibodies against ZO-1 and the nuclear dye DAPI revealed that a subset of cells contained multiple nuclei—typically two—distributed throughout the sheet (**Figs. 1A and 1B**). Further immunofluorescence analysis showed that these multinucleated cells exhibited markedly increased nuclear p53 staining compared to surrounding mononucleated cells (**Fig. 1C**). To test whether multinucleation per se induces p53 activation, we artificially induced multinucleation by treating MDCKII cells with blebbistatin, a myosin II ATPase inhibitor, for overnight. This treatment led to prominent cytokinesis failure, resulting in multinucleation in nearly all cells (**Fig. S1A**). Immunofluorescence staining of these cells revealed elevated nuclear p53 signals in multinucleated cells (**Fig. S1A**). Western blot analysis further demonstrated that p53 protein levels remained significantly elevated in the multinucleated cell population compared to control cells, even 24 hours after blebbistatin washout (**Figs. 1D and 1E**). Notably, the inhibitory effect of blebbistatin on myosin II ATPase activity diminishes approximately 3 hours after washout, indicating that elevated p53 expression observed in this assay is attributable to multinucleation (**Fig. S1B**). These results are consistent with previous reports showing that p53 transcriptional activity increase upon multinucleation^14^, and suggest that p53 upregulation is a general cellular response to multinucleation in epithelial cells.

**Figure 1:**
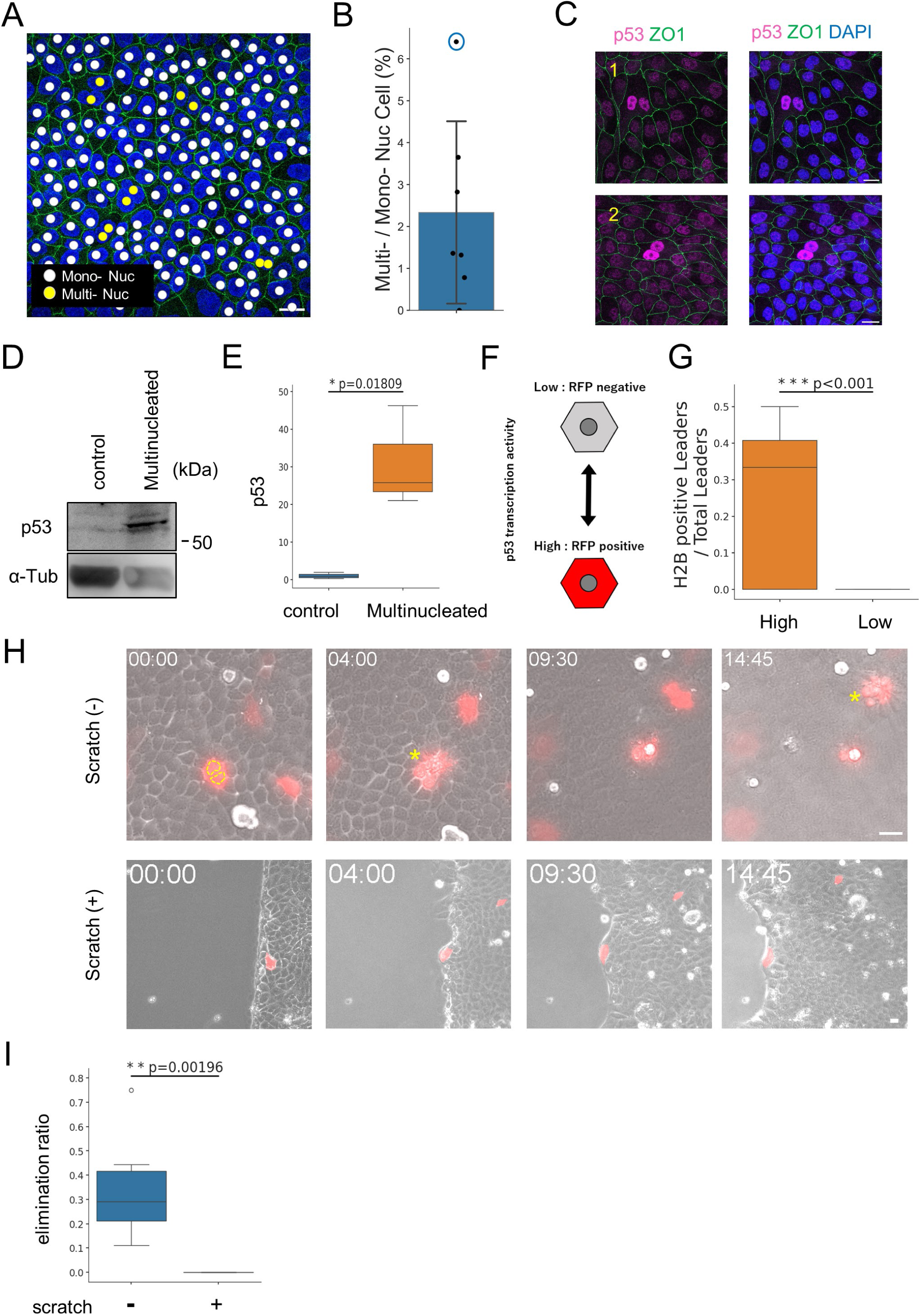
Multinucleated cells scattered within the epithelial sheet exhibit high p53activity and are actively eliminated under normal conditions, but this elimination is suppressed upon epithelial injury. (A) MDCKII cells were seeded and stained for ZO-1 (green) and DAPI (blue). Nuclei of manually identified multinucleated cells are indicated by yellow circles, while those of mononucleated cells are indicated by white circles. (B) Quantification of the percentage of multinucleated cells within the epithelial sheet (n= 7, 1079 Cells). The blue circles indicate the data shown in (A). (C) Representative images of nuclear p53 signal in the MDCKII epithelial sheet. Cells were stained for p53 (magenta), ZO-1 (green), and DAPI (blue). Panels 1 and 2 show images from independent samples. Enhanced nuclear p53 staining was observed in multinucleated cells. (D) Western blot analysis comparing p53 protein levels in control and multinucleated cell populations. Tubulin (α-Tub) was used as a loading control. (E) Quantification of p53 protein levels shown in (D) (n = 3; Student’s t-test). (F) Schematic illustration of the p21 reporter construct. RFP expression occurs in response to the level of p53 transcriptional activity in each cell. (G) Quantification of the elimination ratio of RFP-positive multinucleated cells under low- and high-density conditions using the p21 reporter cell (low: n = 12; high: n = 8; Student’s t-test). (H) Time-lapse imaging of RFP-positive multinucleated cells in MDCKII p21 reporter cells with (+) or without (-) scratch-induced injury. Multinucleation was determined by brightfield imaging (yellow dotted lines indicate nuclei). Yellow asterisks indicate elimination of multinucleated cells from the epithelial sheet. (I) Quantification of the proportion of eliminated RFP-positive multinucleated cells among total RFP-positive multinucleated cells during the observation period (scratch -: n = 6; scratch +: n = 11; Student’s t-test).Scale bars: 20μm.

### Multinucleated cells scattered in epithelial sheets are actively eliminated but this elimination is suppressed upon epithelial injury

It is well established that cells with elevated p53 signaling are eliminated from epithelial sheets due to competitive disadvantage relative to their neighbors, a phenomenon known as cell competition ^20–23^. Based on our observation that p53 expression is markedly elevated in multinucleated cells, we hypothesized that these cells, like those in which the p53 signaling pathway is activated by other factors such as oncogene activation or cellular senescence, may also be subject to elimination from epithelial sheets. To test this, we focused on multinucleated cells with high p53 activity and investigated whether they are indeed eliminated from the epithelial sheet. We introduced a p21 promoter-driven RFP reporter construct into MDCKII cells to monitor cells with elevated p53 transcriptional activity in live-cell imaging (**Fig. 1F**). Time-lapse live-cell imaging revealed that many RFP-positive multinucleated cells—those with high p53 transcriptional activity—were eliminated from the epithelial sheet during the observation period (**Figs. 1G and 1H**, scratch (-)**; Video S1**). These results suggest that multinucleated cells scattered within epithelial sheets are subject to elimination in a manner associated with elevated p53 activity. However, under sparse conditions where the epithelial sheet contains gaps, elimination of multinucleated cells with high p53 activity is suppressed compared to high density conditions (**Fig. 1G**). Based on this observation, we hypothesized that the presence of epithelial gaps, such as those caused by tissue injury, may selectively inhibit the elimination of multinucleated cells.

To test this hypothesis, we used a scratch assay to create artificial wounds in epithelial monolayers and examined the fate of multinucleated cells with activated p53 signaling. Compared to intact epithelial sheets, scratch-induced injury significantly reduced the proportion of RFP-positive multinucleated cells that were eliminated from the sheet during the same observation period (**Figs. 1H and 1I; Videos S1 and S2**). These findings suggest that epithelial injury selectively suppresses the elimination of multinucleated cells with activated p53 signaling.

### Epithelial injury promotes multinucleated cell survival through Akt signaling activation

Given that the suppression of multinucleated cell elimination during epithelial injury was not accompanied by a reduction in p53 transcriptional activity itself (**Fig. 1H**, scratch (+)), we speculated that this phenomenon is not due to a cell-autonomous decrease in p53 signaling. Instead, we considered that survival signals induced by epithelial wounding might play a role in promoting the retention of multinucleated cells.

Akt is a serine/threonine kinase that regulates cell growth and survival. Its activation via phosphorylation, particularly at threonine 308 (T308), promotes cell survival ^24–27^. To assess whether Akt signaling is upregulated upon epithelial injury, we compared the levels of Akt phosphorylation between confluent (normal) and sparsely cultured (injury-mimicking) MDCKII monolayers using Western blot analysis (**Fig. 2A**). This analysis revealed a clear increase in Akt phosphorylation under sparse conditions, with a significant upregulation of the pAkt (T308) signal, suggesting that epithelial wounding triggers Akt activation (**Fig. 2B**).

**Figure 2:**
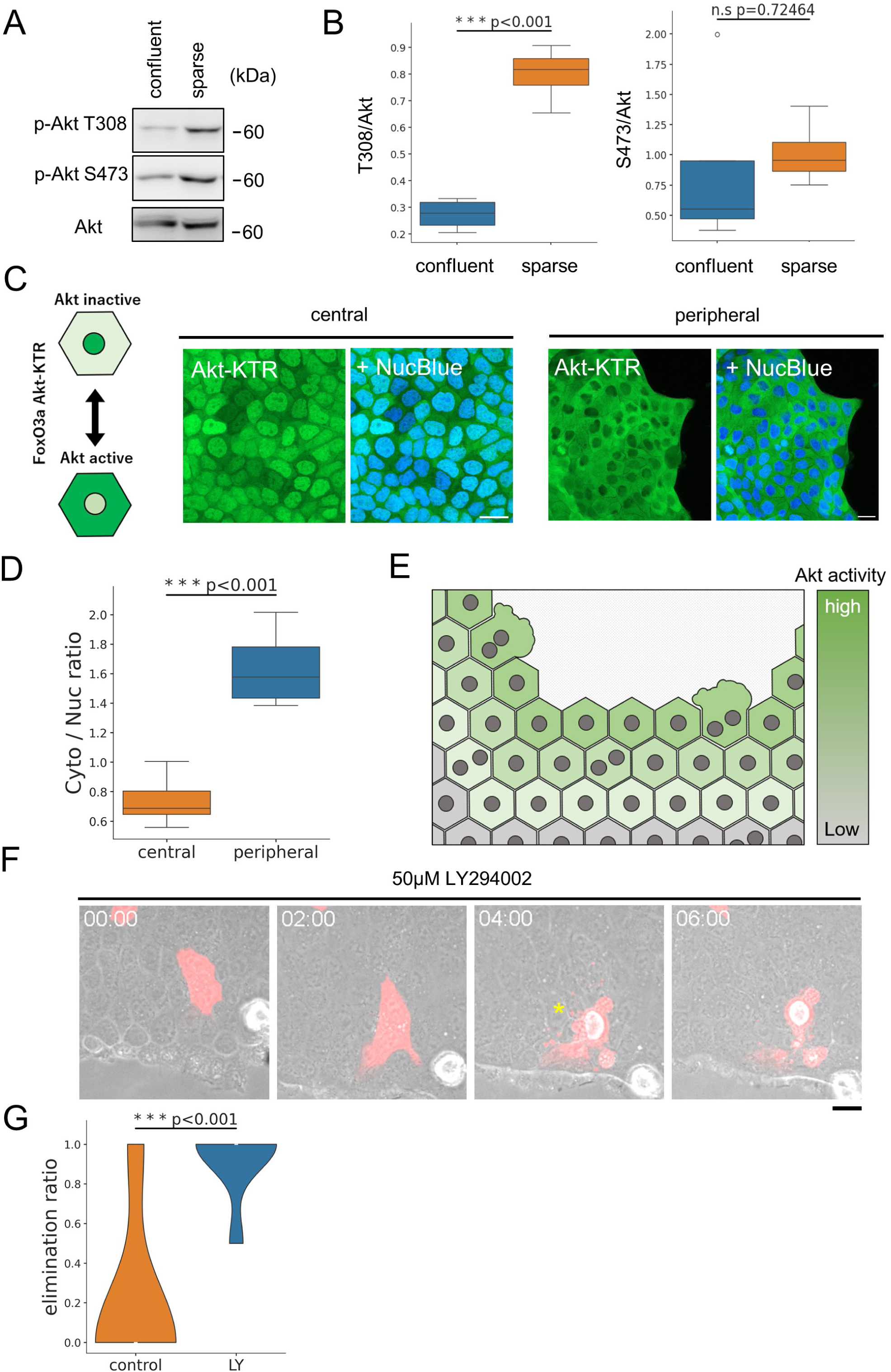
Akt-dependent promotion of multinucleated cell survival occurs upon epithelial sheet injury. (A) Western blot analysis of SDS samples prepared from MDCKII epithelial sheets at confluent and sparse states. Blots were probed for Akt, p-Akt T308, and p-Akt S473. (B) Quantification of the results in (A). Ratios of T308 and S473 signals to total Akt signal are shown (confluent: n = 4, sparse: n = 4; Student’s t-test). (C) Schematic of the FoxO3a Akt-KTR construct and representative images of Akt-KTR signals in central and peripheral regions of the epithelial sheet. NucBlue was added to the medium to label nuclei. (D) Quantification of (C). Ratios of cytoplasmic (Cyto) to nuclear (Nuc) Akt-KTR signal intensities were compared between the central and peripheral regions (central: n = 5, peripheral: n = 4, Student’s t-test). (E) Schematic illustrating the spatial spread of Akt activity upon epithelial injury. Higher Akt activation occurs near the wound edge. (F) Live imaging of p21 reporter cells in the presence of 50μM LY-294002. Multinucleated cells with high p53 transcriptional activity near the wound edge were observed after scratch injury. Yellow asterisk indicates elimination of multinucleated cell from the epithelial sheet. (G) Quantification of (F). The proportion of multinucleated cells with high p53 transcriptional activity that were eliminated was compared between control and LY294002-treated groups (control: n = 6, LY: n = 7; Student’s t-test). Scale bars: 20μm.

To visualize the spatial distribution of Akt activity during injury, we established a MDCKII cell line stably expressing a live-cell Akt activity reporter (FoxO3a Akt-KTR) (**Fig. 2C**)^28^. Live imaging of these cells under wound conditions revealed that cells located away from the wound site displayed predominantly nuclear KTR signal indicative of low Akt activity. In contrast, cells at the wound margin exhibited increased cytoplasmic KTR signal, which reflects enhanced Akt activity near the site of wound (**Figs. 2C and 2D**). These results suggest that epithelial injury induces localized activation of Akt signaling at the wound margin (**Fig. 2E**).

To determine whether this localized Akt activation is required for the survival of multinucleated cells with elevated p53 activity near the wound edge, we treated the monolayers with LY-294002, a PI3K inhibitor that suppresses Akt activation, and monitored the fate of these cells by live imaging (**Figs. S1C and 2F; Video S2**). Under control conditions, p53-high multinucleated cells near the wound site were largely retained within the monolayer (**Figs. 1H and 1I**). However, upon Akt inhibition, these cells were significantly more likely to be eliminated (**Figs. 2F and 2G; Video S2**).

Collectively, these findings indicate that in epithelial cell sheets, tissue injury leads to a local decrease in cell density, which in turn activates the AKT signaling pathway in epithelial cells near the injury site. This injury-induced activation of AKT signaling is required for the survival of p53-high multinucleated cells. Inhibition of AKT activation results in the disappearance of p53-high multinucleated cells from the epithelial sheet, suggesting that without AKT activation, these cells either undergo cell-autonomous apoptosis or are eliminated by neighboring cells. Thus, epithelial wounding appears to confer a survival advantage to these multinucleated cells through the induction of pro-survival signaling, even in the presence of activated p53.

### Multinucleated cells extend protrusions and actively migrate toward epithelial wounds

What is the physiological significance of the prolonged survival of multinucleated cells following epithelial injury? To address this question, we performed time-lapse imaging of p21 reporter cells to monitor the behavior of multinucleated cells located near the wound site. We found that multinucleated cells in the vicinity of the wound extended protrusions toward the wound area. These protrusions originated from multinucleated cells located just inside the wound margin and eventually extended toward the wound substrate (**Fig. 3A; Video S3**).

**Figure 3:**
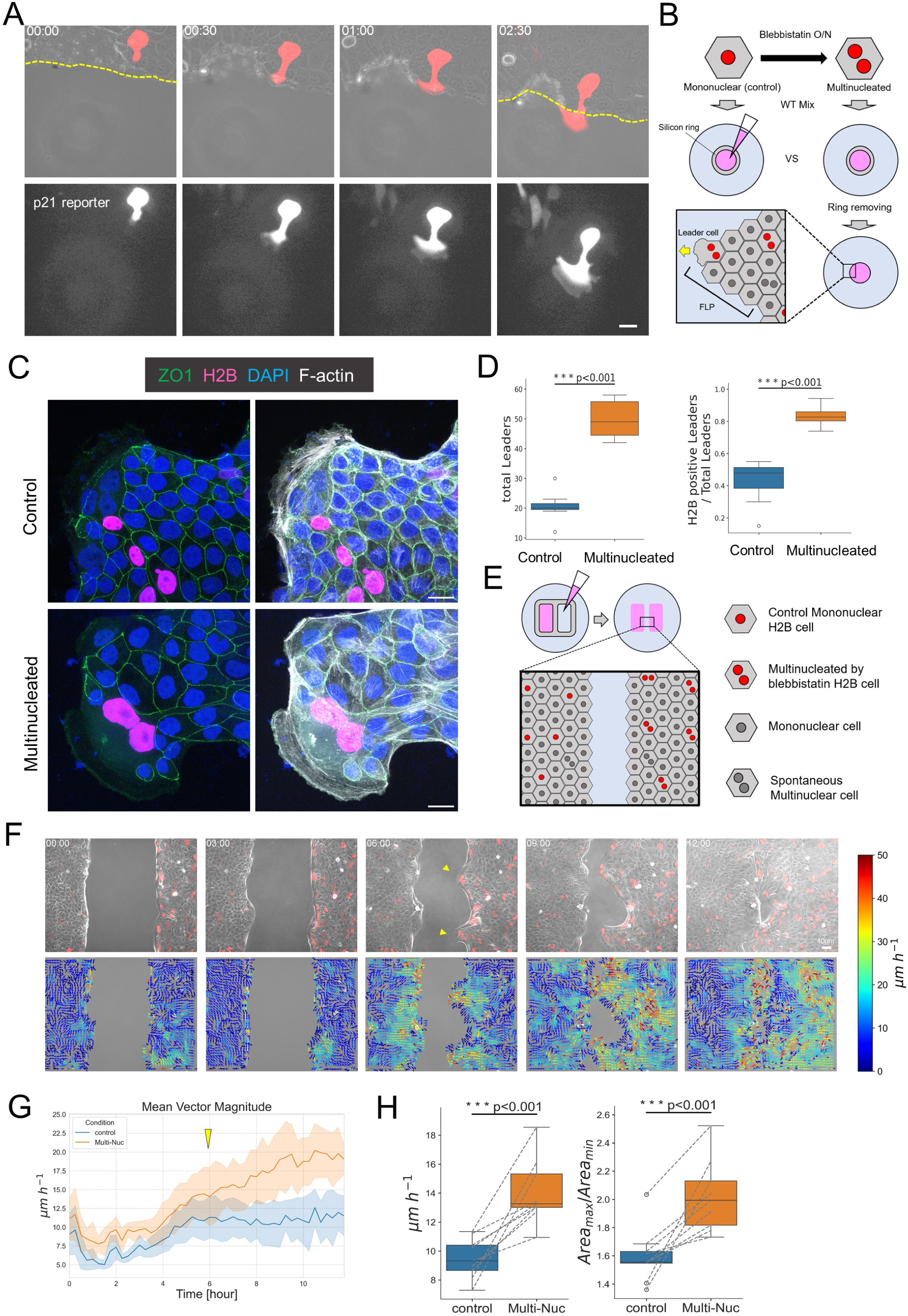
Multinucleated cells preexisting within the epithelial sheet differentiate into leader cells upon tissue injury and enhance wound healing efficiency. (A) Live imaging of p21 reporter multinucleated cells located near the wound edge. RFP-positive multinucleated cells extend protrusions toward the wound area. Lower panels show the RFP signal from the p21 reporter construct in the same field. Yellow dotted lines indicate the boundary between the epithelial sheet and the wound area. (B) Schematic of the experiments shown in (C) and (D). Control (untreated) and multinucleated (induced by overnight blebbistatin treatment) mCherry H2B-expressing cells were mixed with wild-type (WT) cells, seeded into a silicon ring, and cultured until confluency. The ring was then removed to initiate a wound healing assay. (C) Representative images of leader cells at the front of FLP structures formed in the assay shown in (B). Upper panels show the control group; lower panels show the multinucleated group. Cells were stained for ZO-1 (green), DAPI (blue), and phalloidin (F-actin, gray). (D) Quantification of the number of leader cells at FLP tips around the entire wound edge and the proportion of H2B-positive cells among all leader cells (control: n = 7, multinucleated: n = 6; Student’s t-test). (E) Schematic of wound healing assay using culture insert 2-well. Epithelial cells containing untreated H2B-expressing cells were seeded on the left side, and epithelial cells containing multinucleated H2B-expressing cells were seeded on the right side. (F) Representative images during the wound healing assay for control (left side) and multinucleated (right side) groups, with corresponding PIV analysis shown below. Yellow arrowheads indicate leader cells. Scale bars: 40□μm. (G) Time plots of average velocity vectors from PIV analysis in the control (blue) and multinucleated (orange) groups. Yellow arrowheads indicate the point at which epithelial motility increase following leader cell emergence. (H) Quantification of average velocity vectors (left) and wound-covered area (right) (n = 9; paired t-test). Scale bars: 20μm except for (F). Time is indicated as hh:mm.

In addition, immunofluorescence staining of fixed epithelial sheets following scratch injury revealed that these protrusions extended from the basal side of multinucleated cells, traversing beneath adjacent mononucleated cells and reaching the wound substrate (**Figs. S2A and S2B**). These structures were enriched in the focal adhesion components paxillin and FAK^29,30^, suggesting that they are focal adhesion-rich extensions (**Fig. S2C**). Together, these observations indicate that multinucleated cells, once preserved near the wound site, actively form protrusions and crawl toward the injured area, potentially via focal adhesion-mediated motility. Moreover, when the monolayers were fixed and stained one day after scratch injury, finger-like protrusions (FLPs) led by multinucleated cells were observed (**Fig. S2D**). The cells located at the tips of these FLPs are often referred to as “leader cells”, which are known to facilitate collective migration and promote efficient wound closure^3,10^.

These findings suggest that multinucleated cells, once spared from elimination at the wound margin, extend protrusions toward the injury site and may act as leader cells during epithelial repair.

### Multinucleated cells differentiate into leader cells upon epithelial injury and promote wound healing efficiency

Based on the above findings, we hypothesized that multinucleated cells, that preexist within the epithelial sheet prior to injury, differentiate into leader cells upon injury and contribute to wound healing. To test this hypothesis, we examined whether increasing the number of multinucleated cells prior to injury would affect the number of leader cells formed after wounding. Specifically, we established MDCKII cells expressing mCherry H2B to label nuclei and induced multinucleation using blebbistatin (Multinucleated group). Cells left untreated served as the Control group and each treatment population was mixed with the parental wild-type (WT) cells. These mixed cultures were then subjected to a wound healing assay using a silicone ring (**Fig. 3B**; see also Methods section). We found that epithelial sheets containing multinucleated H2B-laebeled cells exhibited a significantly higher number of leader cells compared to controls (**Figs. 3C and 3D**). These results suggest that increasing the number of multinucleated cells prior to wounding leads to a greater number of cells differentiating into leader cells upon injury. Furthermore, the proportion of H2B-positive cells among all leader cells was significantly higher in the Multinucleated group (**Figs. 3C and 3D**), indicating that preexisting multinucleated cells preferentially differentiate into leader cells over mononuclear cells following injury.

Next, we investigated whether the presence of these induced multinucleated cells actually contributes to improved wound healing efficiency. To this end, we seeded epithelial cell populations containing blebbistatin-induced multinucleated H2B cells on the right side of a 2 well culture insert, and control H2B cells on the left side. Once confluency was reached, the insert was removed to initiate a wound healing assay (**Fig. 3E**). The motility of each epithelial sheet was quantitatively assessed using Particle Image Velocimetry (PIV) analysis (**Fig. 3F; Video S4**). The results showed a significant increase in motility in epithelial sheets containing multinucleated cells compared to control sheets (**Figs. 3G and 3H**). In addition, the final area covered by the epithelial sheet was significantly larger in the group containing multinucleated cells induced by treatment, as compared to the untreated control group (**Fig. 3H**). During live imaging, FLPs led by leader cells were frequently observed in the multinucleated group (**Fig. 3F**, 06:00, yellow arrowheads), corresponding with a marked increase in collective cell motility (**Fig. 3G**).

Together, these findings demonstrate that multinucleated cells present in the epithelial sheet prior to injury can function as leader cells upon wounding, thereby enhancing the efficiency of epithelial repair.

### The presence of multinucleated cells is essential for efficient wound healing in epithelial sheets

We next examined whether the presence of multinucleated cells is required for efficient wound healing in epithelial sheets. When MDCKII cells are cultured at low density, they form small colonies, some of which do not contain multinucleated cells. We therefore compared the motility of small colonies with or without multinucleated cells using PIV analysis (**Fig. S3A**). The results revealed that colonies lacking multinucleated cells exhibited significantly reduced motility compared to those containing multinucleated cells (**Figs. S3A and S3B**).

To investigate the requirement for multinucleated cells under more physiological conditions of wound healing, we performed selective depletion of multinucleated cells from larger epithelial cell populations. Since many multinucleated cells exhibit elevated p53 transcriptional activity, they can be identified and removed non-invasively from p21 reporter cells by sorting RFP-positive cells using flow cytometry (**Fig. S3C**). We seeded RFP-negative cell populations (Negative group) on the left side of a 2-well culture insert and unsorted cell populations (Control group) on the right side, followed by removal of the insert to initiate a wound healing assay (**Fig. S3C**; see also Methods section). We found that the Negative group lacking multinucleated cells showed significantly reduced collective motility and decreased wound coverage area compared to the Control group (**Figs. S3D-S3F**).

Together, these results demonstrate that the removal of multinucleated cells from epithelial cell populations impairs wound healing efficiency, suggesting that multinucleated cells play a critical role in promoting epithelial sheet motility during tissue repair.

### Leader cell differentiation and enhanced wound healing driven by multinucleation occur independently of the p53 signaling pathway

Previous study has suggested that upregulation of p53 signaling in leader cells is critical for their function^12^. To determine whether the differentiation of multinucleated cells into leader cells depends on p53 signaling, we generated a p53 knockout (KO) cell line in MDCKII cells (**Fig. 4A**). Interestingly, immunostaining following the wound healing assays using this MDCKII p53 KO cell line revealed that multinucleated leader cells did still form, just like in WT cells (**Fig. 4B**).

**Figure 4:**
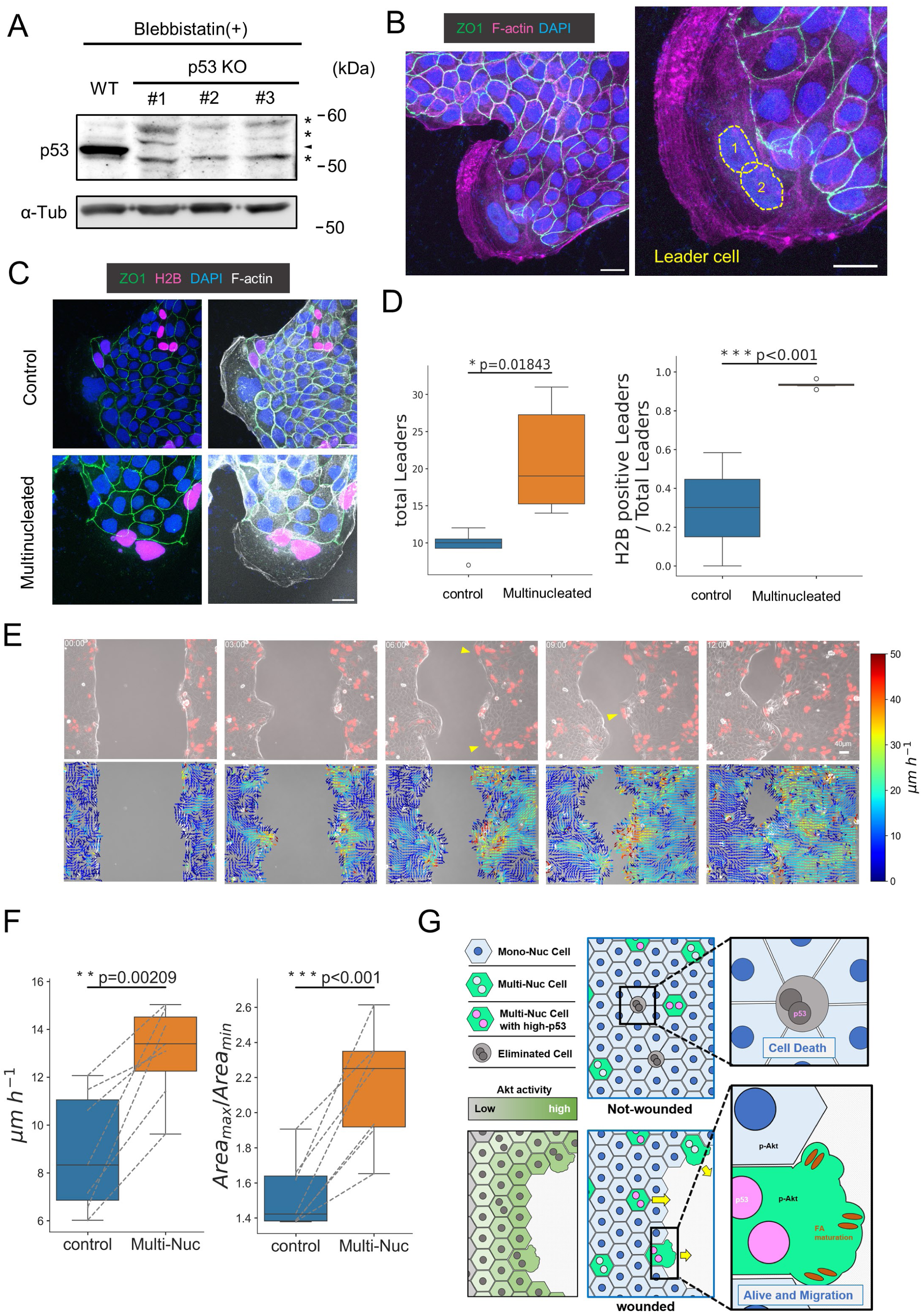
Leader cell differentiation and enhanced wound healing efficiency induced by multinucleation occur independently of p53. (A) Establishment of MDCKII p53 knockout (KO) cell lines. Wild-type (WA) and each CRISPR-edited cell line were treated with blebbistatin overnight and lysed for western blotting against p53. Tubulin (α-Tub) was used as a loading control. Asterisks indicate non-specific bands; arrowheads indicate the p53 band. (B) Formation of FLP structures and leader cell differentiation in the p53 KO cell line. Cells were stained with ZO-1 (green), phalloidin (F-actin, magenta), and DAPI (blue). Yellow dotted lines indicate nuclei. Multinucleation in leader cells is evident. (C) Wound healing assay using the p53 KO cell line as described in Fig. 3B. Representative images of control (top) and multinucleated (bottom) groups. (D) Quantification of the total number of leader cells around the entire wound edge, and the proportion of H2B-positive cells among total leader cells (control: n = 4, multinucleated: n = 6, Student’s t-test). (E) Representative images (top) and corresponding PIV analysis (bottom) of the wound healing assay using control and multinucleated p53 KO cells. Yellow arrowheads indicate leader cells at the FLP tips. Scale bar: 40 μm. Time is indicated as hh:mm. (F) Quantification of average velocity vectors (left) and wound-covered area (right) (n = 7; paired t-test). (G) Graphical summary of this study.

To further examine whether multinucleation-induced leader cell differentiation and enhanced wound healing efficiency are independent of p53, we performed the same experiments described above using the MDCKII p53 KO cells (**Figs. 4C-4F**). Notably, even in the absence of p53, multinucleation led to an increased number of leader cells and significantly enhanced wound healing, comparable to that observed in WT cells (**Figs. 4C-4F**).

These findings demonstrate that the enhancement of leader cell potential and the promotion of epithelial sheet wound healing by multinucleation occur independently of the p53 signaling pathway.

### p53 contributes to the maintenance of multinucleation in leader cells

As described above, enhancement of leader cell differentiation and improved wound healing efficiency following multinucleation occur independently of p53. However, in our experiments, we observed that the total number of leader cells formed during the silicone ring wound healing assay was reduced in p53 knockout (KO) cells compared to WT cells (**Fig. S4A**).

To explain this phenomenon, we hypothesized that the maintenance of multinucleation in leader cells may require p53 signaling, which is known to regulate cell cycle arrest downstream of multinucleation^14^. To test this hypothesis, we examined whether leader cells in p53 KO cells were still proliferative by staining with the proliferation marker ki-67. As expected, leader cells in WT cultures were negative for ki-67 staining, while those in the p53 KO cell line were positive (**Fig. S4B**), indicating cell cycle arrest failure in the absence of p53 in leader cells. Moreover, we observed FLP structures lacking leader cells in the p53 KO monolayers (**Fig. S4C**). This suggests that, without p53, leader cells at the tips of FLPs fail to arrest the cell cycle and undergo cytokinesis, resulting in their division into two mononucleated daughter cells.

Together, these findings suggest that p53-dependent cell cycle arrest is essential for maintaining multinucleation in leader cells, thereby enabling them to sustain their function during epithelial wound healing.

## Discussion

How functionally distinct leader cells emerge from a seemingly homogeneous epithelial sheet upon tissue injury has remained incompletely understood. In recent years, the prevailing model for leader cell emergence during collective cell migration has been the wound-induced specification model. This model proposes that signals arising from tissue injury, such as mechanical stress, TGF-β, and calcium ions, are sensed by cells near the wound edge, which in turn activate signaling pathways involving molecules such as Dll4 and p53 to drive fate commitment toward a leader cell identity^11,12,31,32^. However, this model does not fully explain why only a limited subset of cells among those directly facing the wound differentiate into leader cells.

In this study, we propose a new model of leader cell differentiation that complements the existing wound-induced model. We show that a population of multinucleated cells with high leader potential preexist within the epithelial sheet, and that these cells begin to function as leader cells in response to tissue injury (**Fig. 4G**). This model provides a new perspective on how a small subset of leader cells are selectively emerge upon epithelial wounding.

It was recently proposed that differences in substrate stiffness or topographical features of the wound site can give rise to imbalances in mechanical forces, which in turn contribute to the formation of a limited number of FLPs^1,33–35^. In addition, these mechanical cues are suggested to activate collective cell migration through signaling molecules such as Merlin and ERK^36,37^. Our findings add a new dimension to this concept, suggesting that such mechanical imbalances may not only result from extrinsic factors, but also from intrinsic heterogeneity within the epithelial sheet, specifically, the presence of multinucleated cells. From this viewpoint, investigating whether multinucleated cells possess a heightened capacity to generate mechanical forces could be an intriguing direction for future research.

Multinucleated cells that arise sporadically within epithelial tissues, for example due to cytokinesis failure^17–19^, are traditionally considered abnormal and associated with a pre-neoplastic state^38^. Accordingly, such cells are typically regarded as targets for active elimination in order to maintain epithelial homeostasis^13–16^. However, our study suggests that the survival of multinucleated cells is dynamically promoted in response to tissue injury (**Fig. 4G**). By demonstrating the possibility that the cellular composition of epithelial tissues can be dynamically modulated, this work contributes to a deeper understanding of how epithelial cell populations are organized and functionally diversified.

In this study, we were unable to elucidate the molecular mechanisms underlying why multinucleated cells possess high leader potential. It is well established that leader cells require remodeling of all cytoskeletal components—actin, microtubules and intermediate filaments—as well as interactions with the extracellular matrix to exert their migratory functions^39–42^. Indeed, various cytoskeletal changes were observed in leader cells (**Fig. S5**). However, how multinucleation affects these cytoskeletal networks remains unclear. Previous studies have suggested that amplified centrosomes associated with multinucleation can promote microtubule polymerization and contribute to Rac1 activation. It is therefore possible that Rac1 activation, mediated by enhanced microtubule dynamics, may underlie the increased migratory capacity of multinucleated cells during collective cell migration^14,43^. However, as Rac1 functions at the core of intricate signaling cascades, its precise role in the context of collective cell migration warrants further in-depth study.

Moreover, it remains to be determined whether the enhanced leader potential conferred by multinucleation, as demonstrated in this study, is functionally relevant in vivo. Multinucleation of epithelial cells is reportedly essential for specific physiological functions in various tissues^44,45^. Notably, a recent study showed an increased proportion of multinucleated tubular epithelial cells following acute kidney injury^46^. Based on our findings, it is conceivable that this increase may be partially attributable to enhanced survival mediated by Akt activation and that multinucleated cells may act as leader cells to promote tissue repair following acute kidney injury.

## Materials and methods

### Cell culture

MDCKII and HEK293T cells were cultured in Dulbecco’s modified Eagle medium (DMEM; Nacalai Tesque) supplemented with 10% fetal calf serum (FCS; Sigma-Aldrich). All cell cultures were maintained in a humidified incubator at 37□°C in 5% CO□.

The following primary antibodies were used for immunofluorescence microscopy and immunoblotting: Mouse anti-Paxillin mAb (51-9002034; BD Transduction Laboratories); rabbit anti-p53 pAb, rabbit anti-Akt (pan) mAb (C67E7), rabbit anti-phospho-Akt (Thr308) mAb (C31E5E), rabbit anti-phospho-Akt (Ser473) mAb (D9E), rabbit anti-Myosin IIb pAb, rabbit anti-ki-67 mAb (D3B5), rabbit anti-Vimentin mAb (D21H3; Cell Signaling Technology); mouse anti-cytokeratin 18 mAb (CBL173; LS Bio). Mouse anti-α-tubulin mAb (12G10) and mouse anti-ZO-1 mAb (T8754) were produced in house.

### Plasmids

The p21 reporter plasmid (Addgene #52432) was used to monitor cell cycle arrest. The FoxO3a Akt-KTR construct was kindly provided by Dr. Kazuhiro Aoki (Kyoto University, Japan). For nuclear labeling and cell cycle monitoring, lentiviral constructs encoding H2B, FUCCI, and CRISPR-based p53 knockout were used (dog; 5’-GGCCGGTCCAGGGGCTGTGG-3’).

### Lentiviral Production and Transfection

Transfections were performed using PEI-max (24765-1; Polysciences Inc.). Lentiviruses were produced in HEK293T cells by transfecting the expression vector with the packaging and envelop vectors (psPAX2 and pMD2.G) and harvested by centrifugation. Infections were carried out in low calcium medium with polybrene. MDCKII cells remained in infection media for 48h, followed by antibiotic selection.

Stable MDCKII clones harboring the p21 reporter were generated, and those displaying robust and reproducible RFP induction in response to DNA damage were selected. Clone #6 was used for experiments shown in Fig. 1 and Fig. 2, while Clone #10 was used in Supplemental Fig. 3.

### Scratch assay

MDCKII cells were seeded at 1×10□ cells/mL onto 35-mm glass-bottom dishes (IWAKI) and cultured overnight. Wounds were introduced by scraping the confluent monolayer with a sterile razor blade. Cells were fixed or imaged at designated time points.

To quantify the elimination of p21 reporter–positive cells in Fig. 1 and Fig. 2, epithelial sheets with or without scratch injury were imaged for 18 hours and 6 hours, respectively. At the start of imaging, RFP-positive cells were identified by applying an intensity threshold using ImageJ. Among these, the proportion of cells that underwent apoptosis and were subsequently eliminated from the epithelial sheet during the observation period was calculated.

### Culture-Insert and Ring-Based Wound Healing Assays

For wound-healing migration assays, 2-well culture inserts (ibidi #81176) were used. MDCKII cells were seeded at 1×10□ cells/mL, 70□μL per well. After overnight incubation and confluency, the insert was removed to allow directed migration.

For circular colony formation, a silicone ring (cut with CARL CPN-10 hole punch) was placed on a glass-bottom dish, and MDCKII cells were seeded at 3×10□ cells/mL (20□μL/ring). After confluency was reached, the ring was removed to initiate migration. For co-culture assays with H2B-labeled untreated control or blebbistatin-treated multinucleated cells, cells were mixed with WT MDCKII at a 1:3 ratio (culture insert: WT 1.5×10□/mL + H2B 0.5×10□/mL; silicone ring: WT 4.5×10□/mL + H2B 1.5×10□/mL).

### Cell Sorting

To obtain RFP-negative p21 reporter cells, fluorescence-activated cell sorting (FACS) was performed using the SH800 cell sorter (Sony). Sorted cells were seeded at 3×10□ cells/mL (70□μL/well) into culture inserts for subsequent migration analysis.

Cells reached confluency within 8 hours after seeding, at which point the inserts were removed to initiate migration. Inserts were not kept overnight to minimize the potential reappearance of multinucleated cells that had been excluded during sorting.

### Immunofluorescence

MDCKII cells cultured on coverslips or 35mm glass-base dish were fixed with 0.75 % formaldehyde prepared in PBS for 15 mins at RT and treated with 0.4% Triton X-100 prepared in PBS for 5 mins. Cells were blocked with 1% BSA prepared in PBS for 1h at RT and incubated with primary antibodies for 1 h at RT followed by secondary antibodies for 45 mins at RT. Antibodies were prepared in PBS. Samples were observed at RT with the confocal microscope.

Quantification of multinucleated cells in Fig. 1A was performed using low-magnification images of MDCKII epithelial sheets seeded at 1×10□ cells/mL and stained with ZO-1 and DAPI. The percentage of multinucleated cells was calculated as the number of multinucleated cells divided by the number of mononucleated cells within a single field of view.

### SDS-PAGE and immunoblotting

Samples collected by SDS sample buffer were resolved by SDS-PAGE and transferred to nitrocellulose membranes. After an hour blocking with 5% nonfat milk/0.1% Tween 20/TBS (TBST), membranes were sequentially incubated with primary antibody diluted with 5% nonfat milk/TBST for 1 h at RT or overnight at 4 □ and with HRP-conjugated secondary antibody diluted with 5% nonfat milk/TBST for 45 mins at RT. After each antibody reaction, nitrocellulose membranes were washed five times with TBST for 5 mins. Signals were detected by mixing A (100 mM Tris-HCl [pH 8.5], 0.4 mM p-coumaric acid, 5 mM luminol) and B (100 mM Tris-HCl [pH 8.5], 0.04% H_2_O_2_). Chemiluminescence was captured using the LAS-3000 Imaging system (Fujifilm).

To compare the levels of phosphorylated Akt (T308 and S473) between confluent and sparse epithelial cells, MDCKII WT cells were seeded in 6-well plates at a density of 1 × 10□ cells/mL (confluent) or 2 × 10□ cells/mL (sparse) and cultured overnight. Cells were then lysed in SDS sample buffer for subsequent immunoblot analysis.

### Chemical Inhibitors

To induce multinucleation by inhibiting cytokinesis, blebbistatin (CAY 13186), Myosin II ATPase inhibitor, was used at a final concentration of 50μM and incubated overnight. For the experiments shown in Fig. 2f and 2g, LY-294002 (CAY 70920), PI3K inhibitor, was added to the observation medium (L-15, phenol red-free) at a final concentration of 50μM.

### Live imaging

Time-lapse imaging was performed using either a BZ-X810 (KEYENCE) with 20× or 40× objectives or an LSM900 confocal microscope (Carl Zeiss) with a 63×/1.40 NA oil-immersion objective and a heated stage set at 37□°C. Cells were cultured in Leibovitz’s L-15 Medium, no phenol red (Invitrogen 21083027) supplemented with 10% FCS. Imaging was performed at 15-min intervals for 12–18□h. NucBlue was added to the medium and incubated for 30□min before imaging to enable nuclear segmentation in Akt-KTR-expressing cells.

### Data analysis

Particle image velocimetry (PIV) analysis was performed using a custom Python script based on the OpenPIV^47^ package. For KTR analysis, nuclear segmentation was conducted using Cellpose on NucBlue-stained images. Nuclear and cytoplasmic KTR signals were quantified using nuclear masks. Plots were generated using Seaborn^48^ (Python). Statistical analysis was performed using SciPy^49^: paired t-test (ttest_rel) or Student’s t-test (ttest_ind), as appropriate. All custom scripts used for analysis are available on GitHub [https://github.com/Uta-GitHub/PIVAnalysisForCCM].

## Acknowledgments

We thank all members of the Ikenouchi laboratory (Department of Biochemistry, Graduate School of Medical Sciences, Kyushu University) for helpful discussions.

## Author contributions

Y.T. performed most of the experiments, analyzed the data and wrote the paper. K.M. performed some experiments. J.I. designed research and wrote the paper.

**Supplemental Figure 1:**

**Validation of the inhibitory effects of reagents used in this study**

(A) MDCKII cells were treated with 50 μM blebbistatin and cultured overnight. Three hours after washout, cells were fixed and subjected to immunofluorescence staining with ZO-1 (magenta), p53 (cyan). Multinucleation was observed, and p53 signals were detectable.

(B) The effect of blebbistatin washout. In the “Bleb o/n, 3h after w/o” condition (blebbistatin overnight, 3 hours after washout), actomyosin organization appeared comparable to that of the control, indicating loss of blebbistatin activity. Yellow arrowheads indicate peripheral actomyosin bundles.

(C) Treatment with 50 μM LY294002 for 3 hours (right three lanes) led to the disappearance of p-Akt T308 and p-Akt S473 bands on western blotting.

**Supplemental Figure 2:**

**Characteristics of cellular protrusions extended by multinucleated cells near the wound edge**

(A) MDCKII epithelial sheets were fixed and stained 3 hours after scratch-induced injury. DAPI (blue), phalloidin (F-actin, gray). Yellow dotted lines indicate the boundary between the epithelial sheet and the wound edge. Representative images are shown for the apical, basolateral, and basal planes, corresponding to different Z-slices of the same field of view.

(B) Schematic representation of a multinucleated cell in panel (A). The yellow dotted line outlines the cell body based on apical imaging. The red dotted line marks the protrusive regions identified from the basolateral and basal planes. Nuclei are shown in blue.

(C) Magnified view of the yellow box in (A). Immunostaining for paxillin (green), FAK (magenta), and F-actin (gray) is shown.

(D) MDCKII epithelial sheets were fixed and stained after overnight incubation following scratch injury. ZO-1 (green), DAPI (blue), and phalloidin (F-actin, gray). The yellow boxed region indicates a leader cell located at the tip of a FLP, and is shown at higher magnification in the right panel. Yellow dotted lines indicate nuclear boundaries. The leader cell is confirmed to be multinucleated. Scale bars: 20μm.

**Supplemental Figure 3:**

**Multinucleated cells are essential for efficient wound healing of epithelial sheets**

(A) Time-lapse imaging and PIV analysis of small MDCKII epithelial colonies. mCherry H2B-expressing cells indicate multinucleated cells.

(B) Quantification of colony motility from (A). Comparison between mononuclear (Mono-) and multinucleated (Multi-Nuc) colonies. (Mono-: n = 3, Multi-Nuc: n = 3, Student’s t-test).

(C) Schematic of the experiment shown in (D–F). p21 reporter cells were sorted using a cell sorter to remove RFP-positive (multinucleated) cells, creating a “Negative” group. Middle panel shows fluorescence microscopy of RFP-positive cells; yellow numbers indicate nuclear counts. The representative RFP-positive cell has two nuclei. Cells were seeded in a culture insert (2-well), with Negative cells on the left and unsorted control cells on the right, followed by wound healing assay.

(D) Upper panels: time-lapse images from the experiment in (C). Lower panels: corresponding PIV analysis. Scale bars: 40 μm. Time is indicated as hh:mm.

(E) Time-course plots of average vector magnitude obtained from the PIV analysis in, comparing Negative (blue) and Control (orange) groups (n = 4 each).

(F) Quantification of average velocity vectors (left) and the area covered by the epithelial sheet (right) (n = 4, paired t-test).

**Supplemental Figure 4:**

**p53-dependent cell cycle arrest in leader cells is required for the maintenance of multinucleation**

(A) Comparison of the total number of leader cells between WT and p53 KO cell lines. (WT: n = 7, KO: n = 4, Student’s t*-*test).

(B) Immunostaining for Ki-67 (green) in leader cells from WT and p53 KO cells. In WT cells, leader cells are Ki-67-negative (white arrowhead), indicating cell cycle arrest. In contrast, leader cells in p53 KO cells are Ki-67-positive (red arrowhead), suggesting continued cell cycle progression. Phalloidin (F-actin, gray); DAPI (blue).

(C) FLP structures in p53 KO cells lacking leader cells. ZO-1 (green), phalloidin (F-actin, magenta), DAPI (blue). Scale bars: 20μm.

**Supplemental Figure 5:**

**Cytoskeletal remodeling in leader cells**

(A) Immunostaining of FLP with leader cells at the front. Right panel shows a magnified merged image. Phalloidin (F-actin, magenta), α-Tubulin (green).

(B) Immunostaining of FLP with leader cells at the front. Bottom panels show magnified views of boxed regions in the merged image: yellow box 1 (leader cell) and yellow box 2 (follower cell). CK18 (magenta), vimentin (green). Scale bars: 20μm.

