## Supplementary figures and images for "Wounding induces preexisting multinucleated cells to survive and thrive as leader cells"

### Figure S1

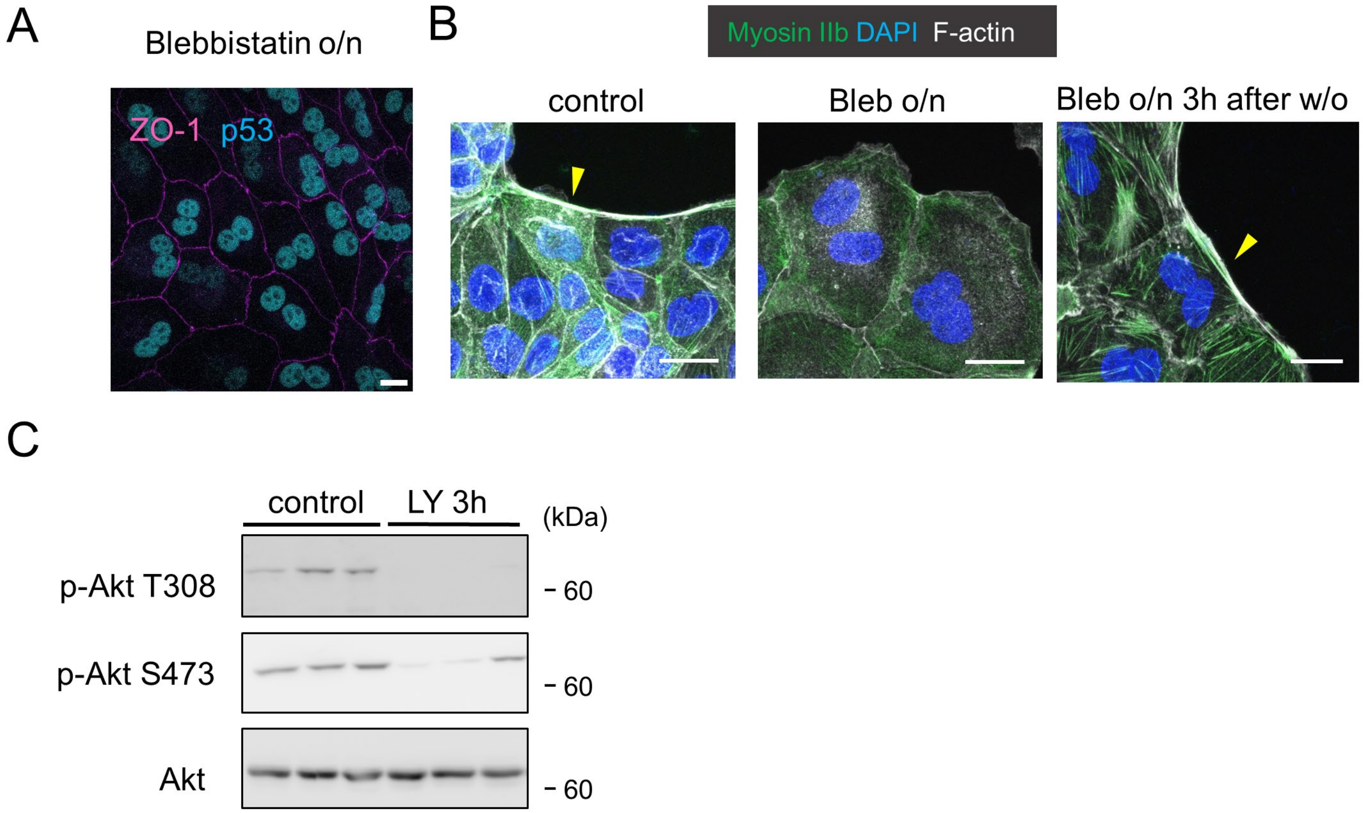


Supplemental Figure 1 Takahashi *et al*.

### Figure S2

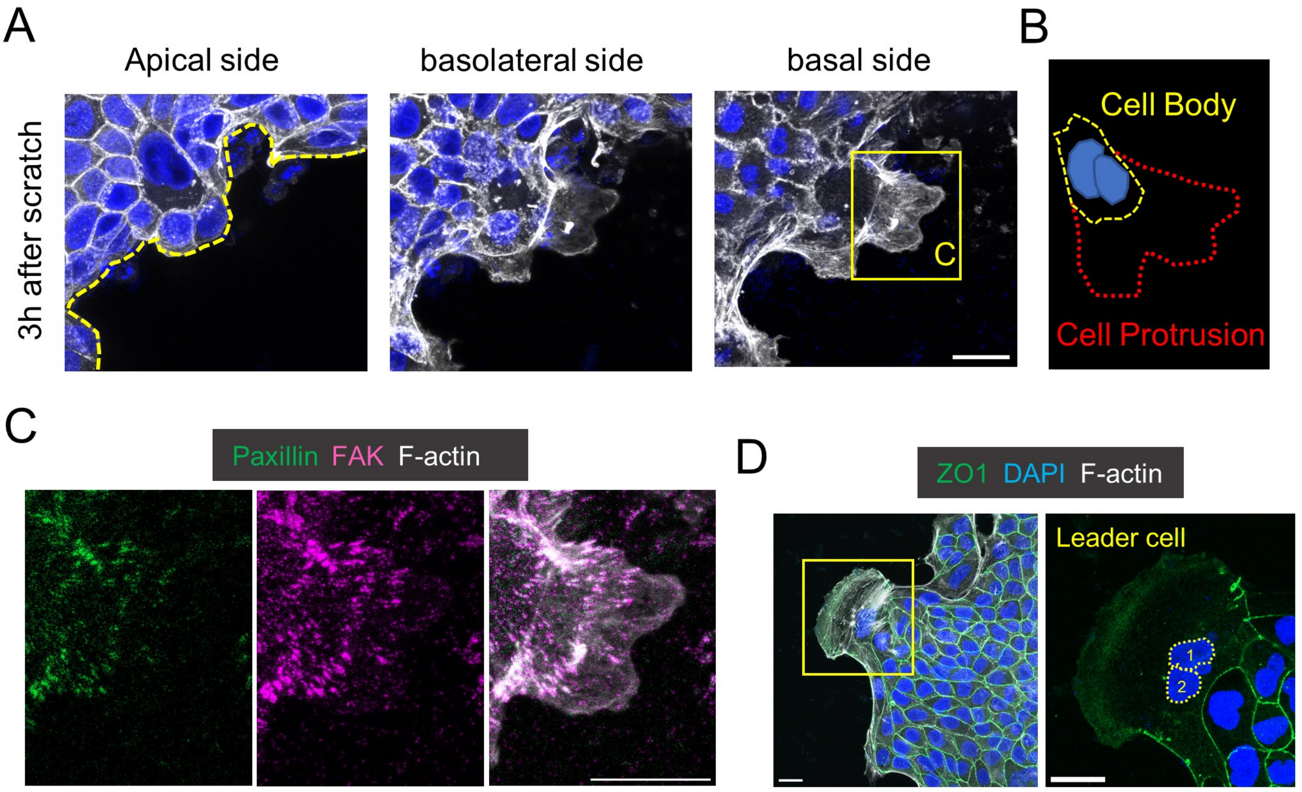


Supplemental Figure 2 Takahashi *et al*.

### Figure S3

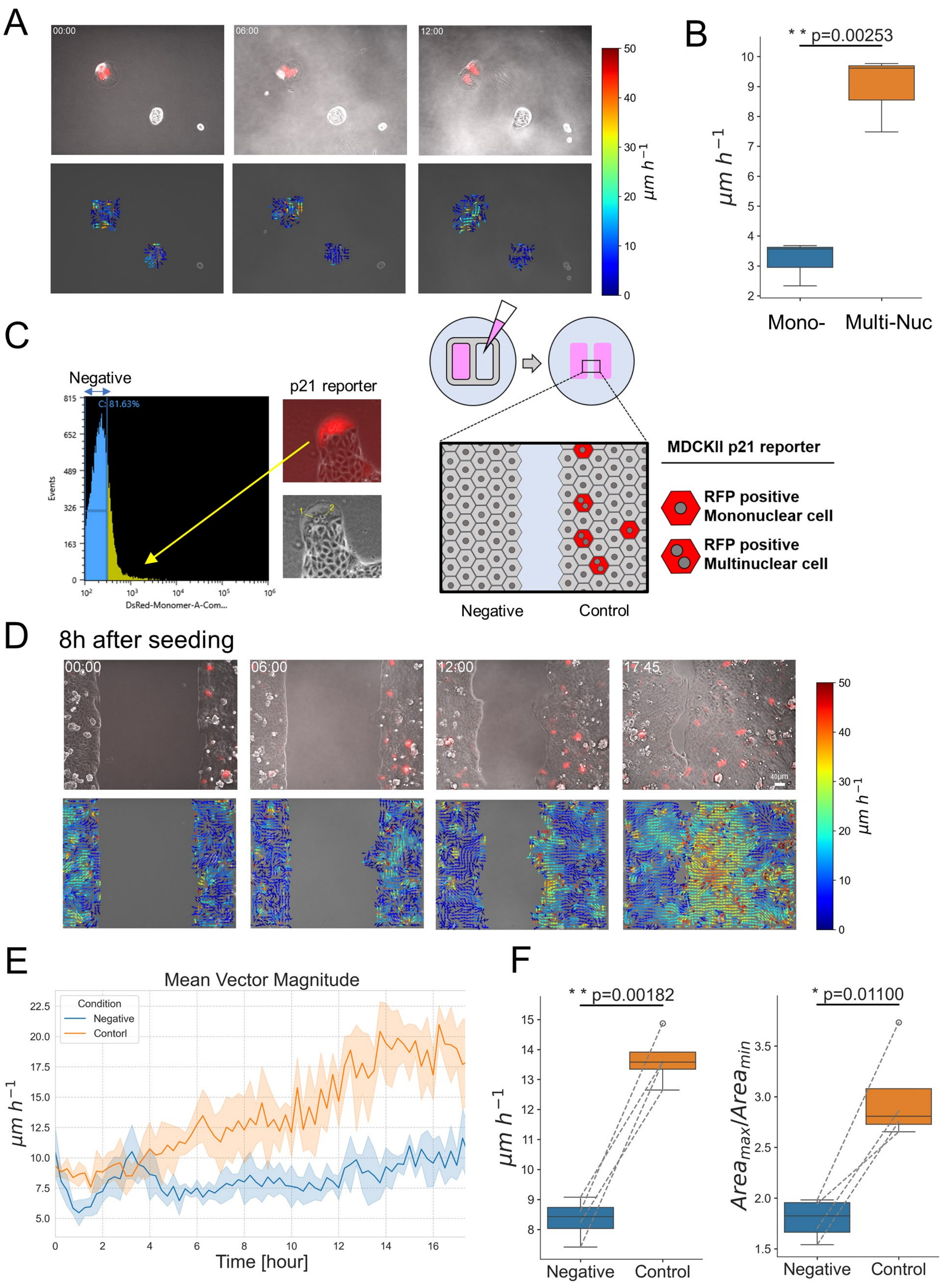


Supplemental Figure 3 Takahashi *et al*.

### Figure S4

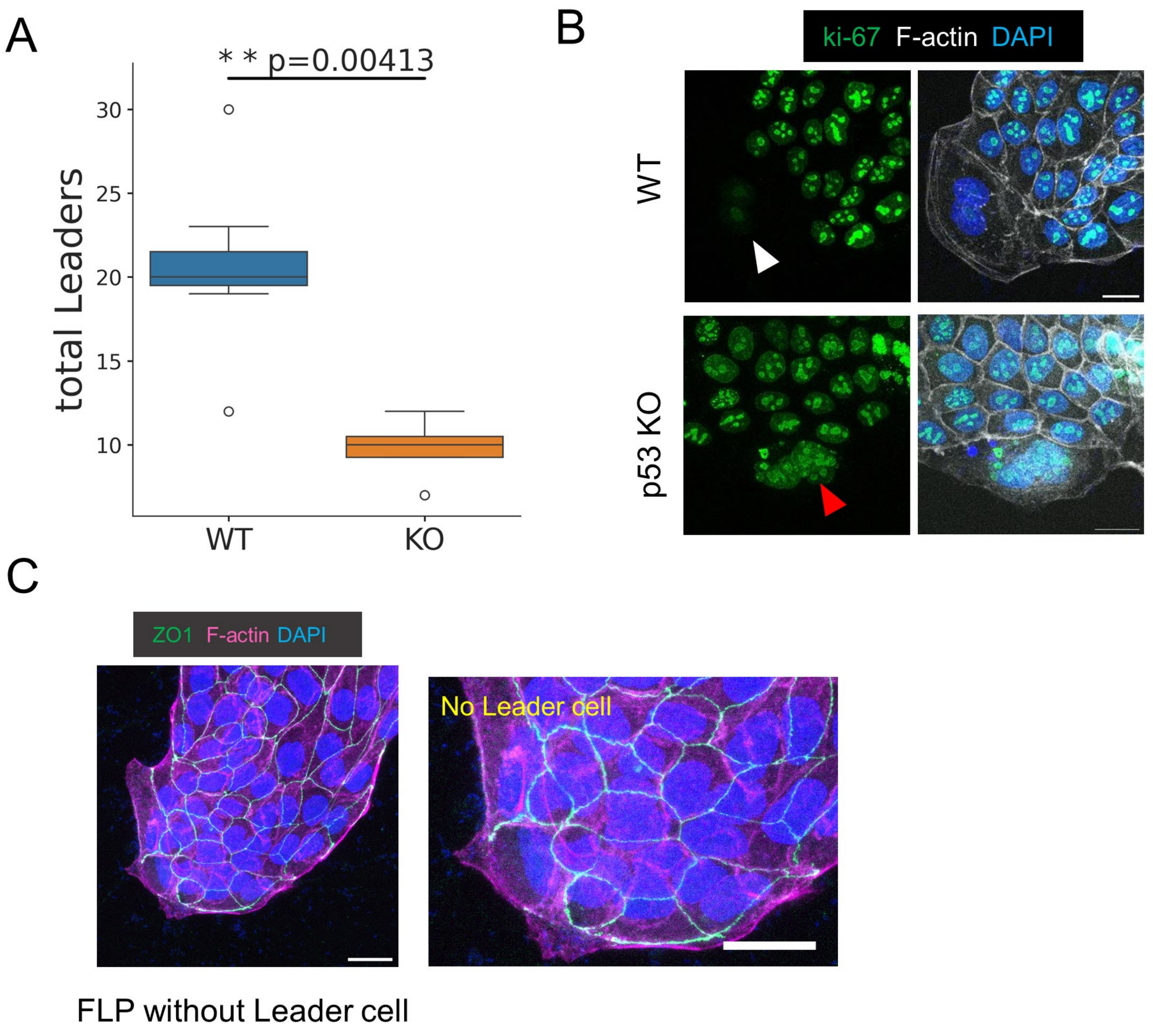


Supplemental Figure 4 Takahashi *et al*.

### Figure S5

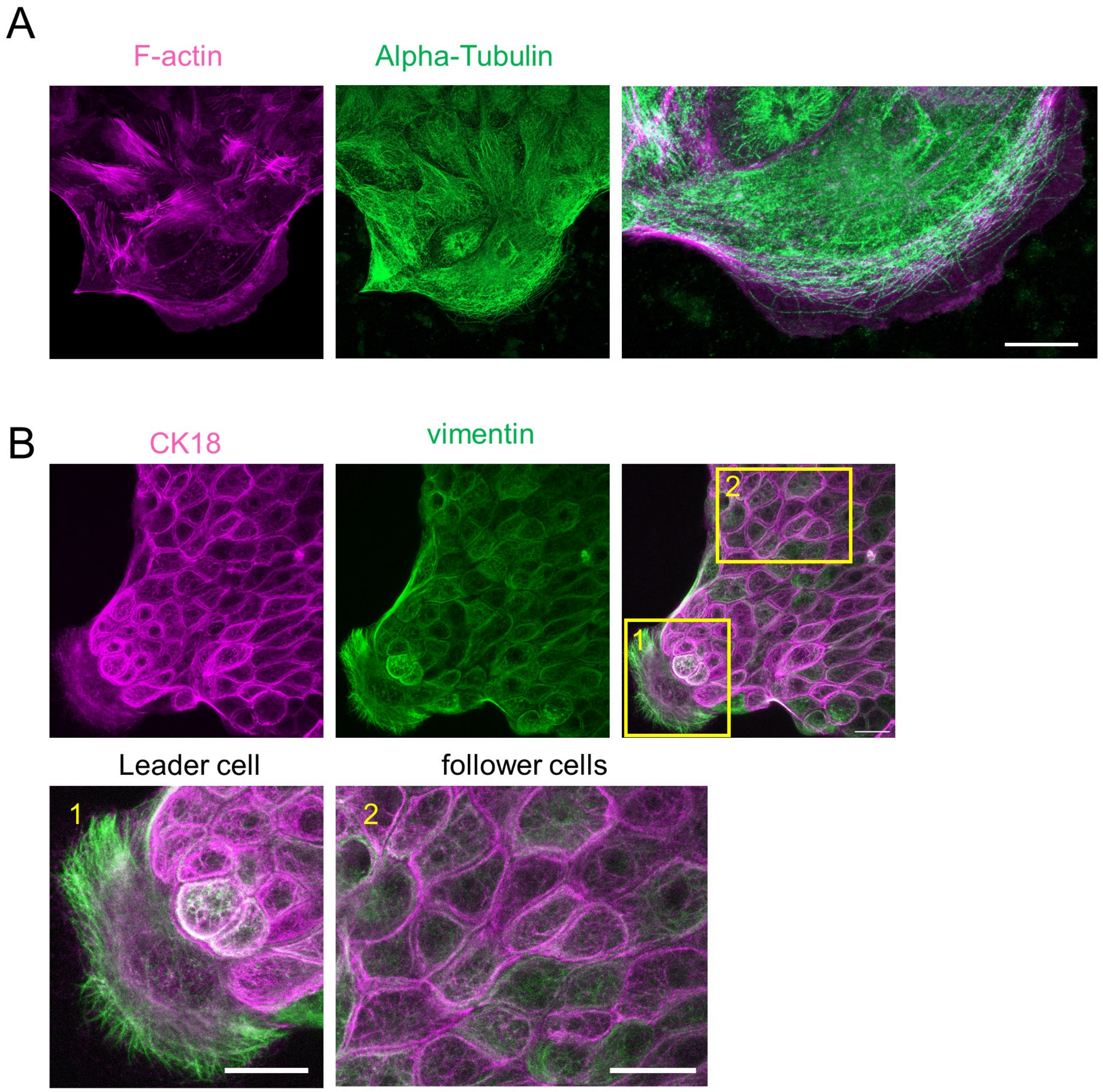


Supplemental Figure 5 Takahashi *et al*.
